# Cheating the cheater: Suppressing false positive enrichment during biosensor-guided biocatalyst engineering

**DOI:** 10.1101/2021.10.08.463720

**Authors:** Vikas D. Trivedi, Karishma Mohan, Todd C. Chappell, Zachary J. S. Mays, Nikhil U. Nair

## Abstract

Transcription factor (TF)-based biosensors are very desirable reagents for high-throughput enzyme and strain engineering campaigns. Despite their potential, they are often difficult to deploy effectively as the small molecules being detected can leak out of high-producer cells, into low-producer cells, and activate the biosensor therein. This crosstalk leads to the overrepresentation of false positive/cheater cells in the enriched population. While the host cell can be engineered to minimize crosstalk (e.g., by deleting responsible transporters), this is not easily applicable to all molecules of interest, particularly those that can diffuse passively. One such biosensor recently reported for *trans*-cinnamic acid (tCA) suffers from crosstalk when used for phenylalanine ammonia-lyase (PAL) enzyme engineering by directed evolution. We report that desensitizing the biosensor (i.e., *increasing* the limit of detection, LOD) suppresses cheater population enrichment. Further we show that, if we couple the biosensor-based screen with an orthogonal pre-screen that eliminates a large fraction of true negatives, we can successfully reduce the cheater population during the fluorescence-activated cell sorting (FACS). Using the approach developed here, we were successfully able to isolate PAL variants with ~70% high k_cat_ after a single sort. These mutants have tremendous potential in Phenylketonuria (PKU) treatment and flavonoid production.

## INTRODUCTION

While experimental and computational pathway along with biocatalyst designs have made major strides in past decades, screening for high target metabolite producing cells and enzymes remains a major bottleneck^1^. As with protein directed evolution, one of the bottlenecks in combinatorial metabolic engineering is the screening step^2^. Techniques like high-performance liquid chromatography (HPLC) that are traditionally used monitor concentrations of products or metabolic intermediates during strain development are too low throughput to screen large combinatorial libraries^3^. Resultantly, significant effort has been expended to develop higher throughput screening methods to monitor metabolite levels^4–6^. Most popular among these are genetically-encoded biosensors^7–9^, which link a phenotype to a readily detectable quantitative output signal. Biosensors are most frequently transcription factor- (TF) or riboswitch-based^10–12^ – although other modalities like enzyme-based^13–15^ or protein-protein interaction-based^16^ are also possible. Among these, TF-based biosensors are most widely used due to ease of construction and tunability. These TFs specifically bind a target metabolite (e.g., pathway intermediate, substrate, or final product) and activate or repress expression of a reporter gene – usually a fluorescent protein, which can then be screened using fluorescence-activated cell sorting (FACS). Many of the biosensors described in the literature are based on natural transcription factors. However, genetic circuits controlling them often need to be engineered to reduce high basal expression, improve dynamic range, increase sensitivity, and ensure orthogonality^17^. Examples for the application of these TF-based sensor-selector systems include screening campaigns for identifying improved producers of malonyl-CoA^18, 19^, naringenin^20, 21^, *cis,cis-muconic* acid^22, 23^, glucaric acid^24^, and fatty acyl-CoA^25^, vanillin^26^, protocatechuate^27^, butanol^28^, lactam^5^, etc.

A major shortcoming of many biosensor-based systems is the propensity of “cheater” cells or false positives to enrich^1, 29–31^. This is particularly concerning when the metabolite being sensed can be actively or passively transported in and out of cells. Recent examples include use of a biosensor-based engineering to identify overproducers of *trans-cinnamic* acid (tCA)^32^, benzoic acid and derivates^33^, 1-butanol^34^, and naringenin^2^, or to improve synthetic methylotrophy with formaldehyde-responsive TFs^35, 36^. All these studies had to contend with a false positive when using the biosensor for strain or protein engineering. Flachbart et al.^32^ recognized the high false rate and attempted to mitigate it by maintaining the cells at low cell densities to reduce extracellular tCA concentrations and transport. However, their directed evolution campaign to identify improved variants of enzyme phenylalanine ammonia-lyase (PAL) was still severely hindered by presence of false positive cells (“cheaters”) in the FACS-enriched population, resulting in variants with only modest improvement (≤11 %) in k_cat_ even after five rounds of stringent sorting. In this work, we describe conditions that help mitigate enrichment of cheaters during high-throughput screening campaigns using the tCA biosensor system. We first demonstrate that decreasing sensitivity (i.e., increasing the limit of detection, LOD) of the biosensor circuit output significantly reduces false positives. Next, we show that incorporation of a pre-screen can further mitigate cheater enrichment. With the modified workflow, we undertook a directed evolution campaign to improve PAL (from *Anabaena variabilis*^37^) activity on its native substrate, phenylalanine (Phe), and were able to identify variants with ~70 % higher activity (k_cat_) after a single round of sorting – the highest reported for a PAL by biosensor-guided engineering.

## RESULTS AND DISCUSSION

### Abundance of cheater cells and true positives are highly correlated

The *E. coli* HcaR transcription factor (TF), which induces the expression of the hydroxycinnamic acid (*hca*) catabolic operon, forms the basis of our biosensor design, and is similar to the previous design^32^ (**Figure 1A**). We placed *sfGFP* under the control of an HcaR-responsive promoter (P_*hcaE*_) in a plasmid (pVDT46) and transformed in *E. coli* MG1655 *rph^+^* to create a strain hereon referred to as Ecvdt46. However, unlike in the published design, we did not knockout the native *hcaREFCABD* operon in *E. coli,* enabling catabolism of tCA^38^. To assess the evolution of cheater cells by tCA cross-feeding, we investigated how they evolve as a function of true positive population. We call true positives biosensor-encoding cells that generate tCA through PAL-mediated deamination of phenylalanine (Phe) (i.e., PAL^+^) (**Figure 1B**) and true negatives as those without functional PAL (i.e., PAL^−^). We integrated a blue fluorescent protein (BFP) reporter in PAL^−^ cells to aid in tracking their population (**Figure 1C**). Thus, PAL^−^ cells are unable to generate intracellular tCA (true negatives, **Figure 1C)** and would only activate their biosensor if they import exogenous tCA (false positives, **Figure 1D**). Next, we generated a mock library by mixing PAL^−^ and PAL^+^ cells in different ratios and monitored the evolution of cheater cells during co-culture. We observed that the abundance of cheater cells was positively and monotonically PAL^+^ dose-dependent (**Figure 1E-G**). These cells show up in the GFP^+^ and BFP^+^ channels (top right) whereas true positives are GFP^+^ only and are simply upshifted (top left). This validates our hypothesis that false positives are generated only in the presence of PAL^+^ (tCA-producing) cells through import of tCA in PAL^−^ cells. Thus, the system as is, unsuitable for biocatalyst engineering campaign.

**Figure 1.**
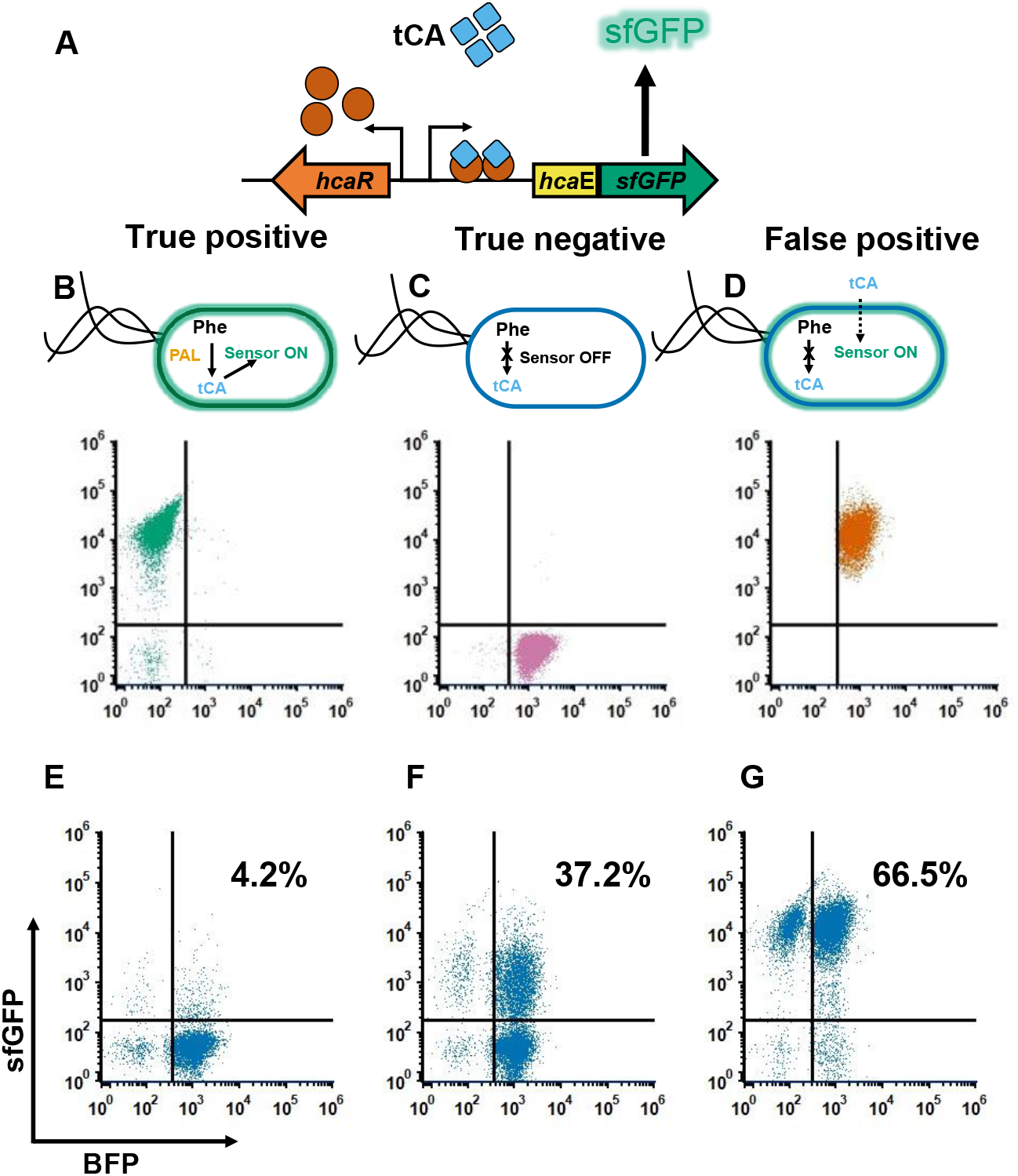
Quantification of cheater cell evolution using flow cytometry. **A)** Schematic representation of tCA biosensor. In the presence of tCA, HcaR servers as the transcriptional activator of *P_hcaE_-sfGFP*. **B-D)** Scatter plots of the control population, PAL^+^ (green, upper left quadrant) and PAL^−^ (pink, lower right quadrant) and cheater cells (orange, upper right quadrant). **E-G)** Mock libraries containing PAL^+^ and PAL^−^ in ratios of 1:100, 1:10, 1:1, respectively. The percentage of cheater cells (upper right quadrant) increased from 4.2% to 66.5% with increasing relative abundance of true positives.

Various approaches have been used to fine-tune the biosensor response, increase their dynamic range, sensitivity, and prevent crosstalk^17^. But these approaches are generally designed and tested on a clonal population, often with exogenously added metabolite, which may not always translate when applied to a high-throughput screen of a heterogenous population like a biocatalyst library with intracellularly produced metabolite. One approach that seems promising is to alter the expression of the TF^39^ (HcaR, here) or titrating the operator binding sites on promoter driving expression of the reporter protein^40, 41^ (in this case, P_*hcaE*_). However, this can be laborious and/or time-consuming. So, we sought an alternate approach for the same outcomes by leveraging the native *E. coli* regulatory mechanism, given that the biosensor-promoter pair is native to this bacterium.

### Carbon catabolite repression (CCR) alters the sensor response to suppress cheater cell evolution

HcaR is an activator of *hca* operon^38^, which encodes genes required to catabolize phenolic acids like tCA. But as non-preferred substrates, the expression of the *hca* operon (and HcaR) is subject to carbon catabolite repression (CCR) by glucose^42^. We hypothesized that this CCR could be leveraged to desensitize the biosensor to low intracellular tCA concentrations present in cheater cells that import exogenous tCA (**Figure 2A**). This creates a threshold for activation that may only be surpassed at high intracellular tCA levels, as expected of cells with active PAL. To test this, we re-created a mock a library of PAL^+^ and BFP-tagged PAL^−^ biosensor cells, as before, and looked for the appearance of cheater cells under different growth conditions – in LB, LB + Phe, glycerol (Gly) + Phe, or glucose (Glc) + Phe. We observed, as expected, that in non-repressing media (LB, LB + Phe, Gly + Phe) the percent of cheater cells was high (**Figure 2B-D**), and *vice versa* in Glc + Phe condition (**Figure 2E**). This supports our hypothesis that native regulatory structure can be leveraged to suppress cheater evolution. To further support our conclusion that Glc-mediate CCR desensitizes the biosensor, enabling activation only at higher intracellular concentrations, we assessed the response of the biosensor to exogenously added tCA in both, Glc and Gly, media (**Figure 2G**). Although the fluorescence response for both Gly and Glc containing media was similar (**Figure S2A**), Glc condition exhibited higher fold change in fluorescence (150-fold) and larger dynamic range (100 – 750 μM, **Figure 2H**). We also observed activation of the tCA sensor in media containing Gly even when not challenged with tCA (**Figure S2B**). Whereas we observed drop in fluorescence in Glc containing media (**Figure S2B**). The EC_50_ observed for Glc (386 μM) condition is higher than that for Gly (105 μM), indicating decreased sensitivity. This decreased sensitivity appears to be beneficial as it suppresses fluorescence activation in cheater cells that are expected to have a lower intracellular concentration of tCA.

**Figure 2.**
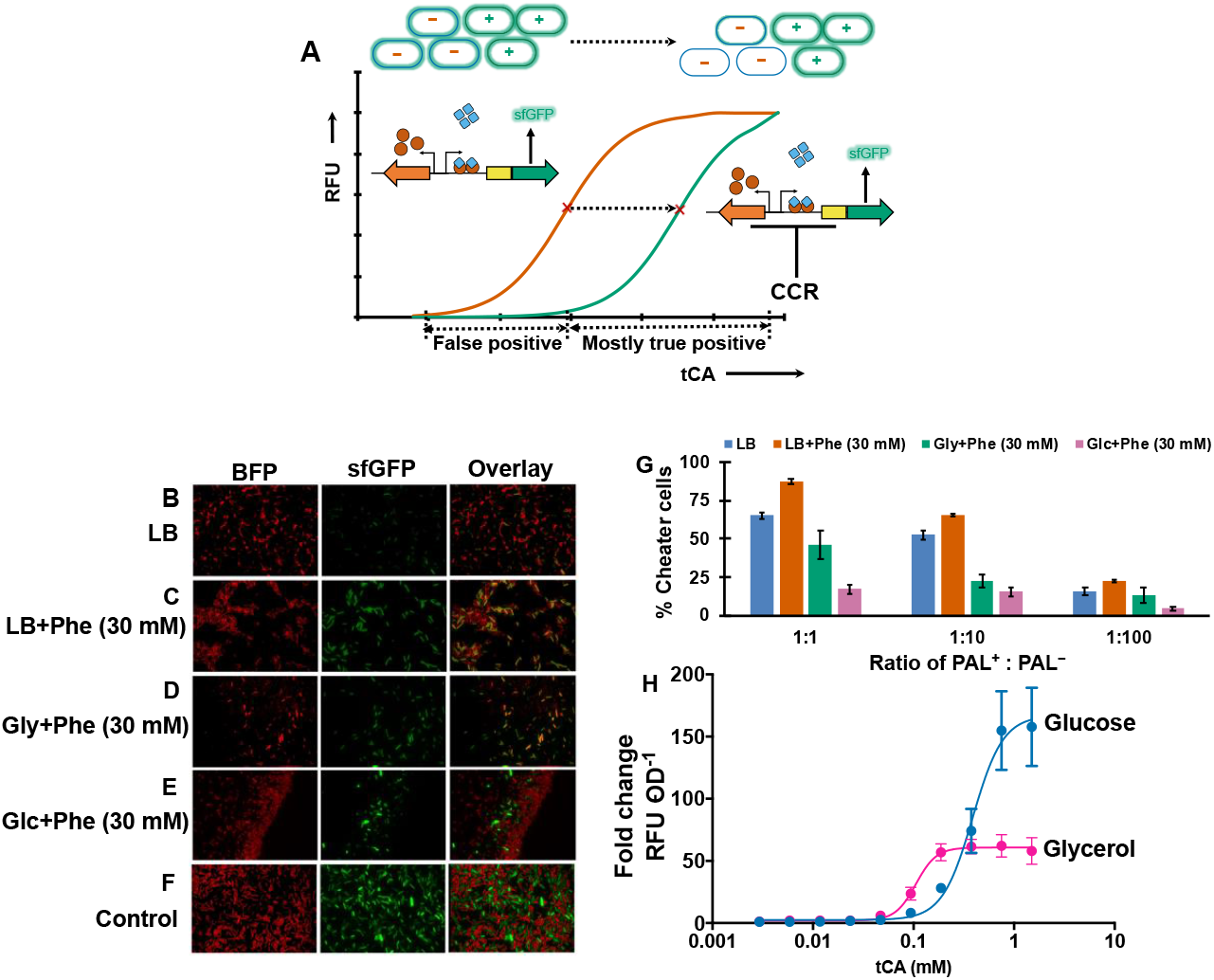
Evaluation of cheater cell evolution under CCR. **A)** Schematic of how CCR can be exploited to suppress cheater cells by right-shifting the EC_50_ of the synthetic circuit. PAL^−^ cells (blue outline), cheater cells (green glow with blue outline), A co-culture of PAL^+^ and BFP-tagged PAL^−^ biosensor strain cells were mixed in 1:1 ratio and grown for 8 h before imaging. Mixture grown in **B)** LB, **C)** LB + Phe (30 mM), **D)** Gly + Phe (30 mM), and **E)** Glc + Phe (30 mM). True positives are green (sfGFP), true negatives red (false-colored BFP). Cheater cells are yellow (red + green). **F)** Control – cells not co-cultured together but mixed immediately before imaging. **G)** Percent of cheater cells as determined from flow cytometry for the different growth conditions. The flow cytometry profiles are given supplemental section (**Figure S1, Table S1**). **H)** The dynamic range of fluorescent output in glucose or glycerol when supplemented with tCA (3 μM – 3 mM) after 8 h of growth in complete minimal media. The fluorescence output normalized to uninduced control of the respective carbon sources.

### An orthogonal growth-coupled pre-screen further suppresses cheater evolution

Previously, we developed and optimized an enrichment for active PAL variants by linking evolution of ammonium (NH4^+^) to cell growth during deamination of Phe to tCA^37^. We also recently showed that this screen readily and rapidly eliminates inactive PAL variants from a mutagenized library^43^. Hence, we posited that using this orthogonal pre-screen based on growth can further mitigate cheater cell enrichment during FACS (**Figure 3**). For this, we created a mock library containing PAL^+^ and BFP-tagged PAL^−^, as before (in 1:1 and 1:10 ratios, respectively). This pre-defined library was subjected to growth-coupled enrichment for three passages in selective minimal medium with Phe as the sole nitrogen source (**Figure 3A**). At the end of every passage, the samples were analyzed on flow cytometry to detect the presence of cheater cells (**Figure 3B–E**). We observed a drop in the percent of cheater cells in the population and at the end of passage #3 to <1 % (**Figure 3F**). This indicates that pre-screening of library using growth-coupled enrichment can largely eliminate the PAL^−^ population and reduce the number of cheater cell events during FACS (**Figure 3G**).

**Figure 3.**
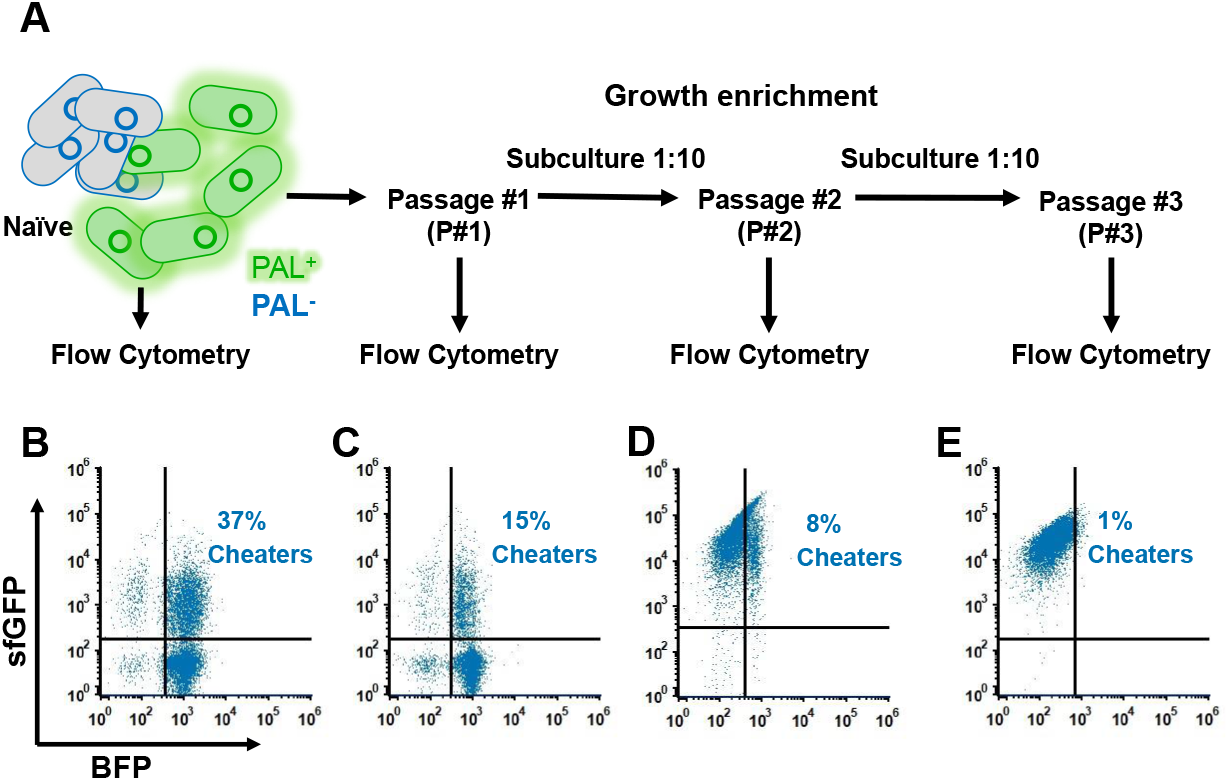
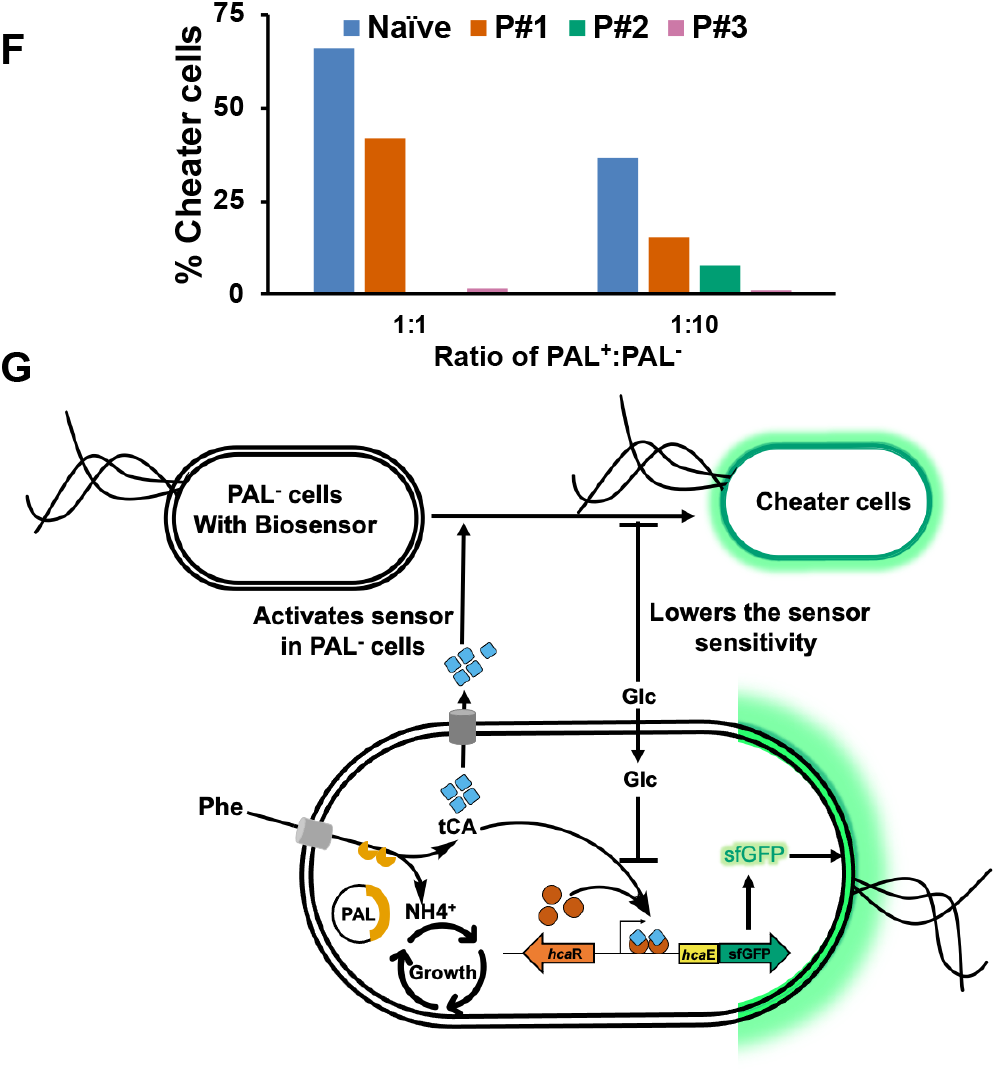
Growth-coupled enrichment mitigates cheater evolution. **A)** Overview of method used to demonstrate that growth-coupled pre-screen eliminates cheater cells. PAL^+^ and BFP-tagged PAL^−^ cells were mixed in 1:1 or 1:10 ratio and subjected to selection for three passages in selective media. Presence of cheater cells were shown for 1:10 mock library using flow cytometry for **B)** Naïve, **C)** passage #1 (P#1), **D)** passage #2 (P#2), and **E)** passage #3 (P#3, gating was adjusted for P#2 and #3 to account for increased sfGFP fluorescence which could not be corrected by compensation). See supplemental **Figure S3** for scatter plots of the 1:1 mock library. **F)** Percent of cheater cells after different passages relative to naïve for 1:1 and 1:10 mixtures. **G)** Proposed design to suppress activation of the biosensor in cheater cells through carbon catabolite repression and growth-coupled pre-screen. In presence of active PAL, intracellular phenylalanine (Phe) is deaminated to ammonium (NH4^+^) and *trans*-cinnamic acid (tCA). tCA binds to HcaR to activate the transcription of *P_hcaE_-sfGFP.* Since tCA can be transported in and out of cells, it can enter PAL^−^ cells (that inherently do not produce ay tCA) and activate expression of sfGFP. These PAL^−^ cells show up as “cheater” cells during FACS and flow cytometry. While glycerol (Gly) and glucose (Glc) can serve as substrates for growth, only Glc engages CCR of the *hca* operon, decreasing basal activation. CCR also lowers the sensitivity of the biosensor, which is beneficial in preventing activation of the sensor by low intracellular tCA concentrations that may be present in PAL^−^ cells. Utilization of NH4^+^ for growth produced by PAL can further enrich true positives over false positives in mixed populations.

### The biosensor response is a predictor of enzyme activity

Using the two-step approach, we screened a previously created ~10^5^ member error-prone PCR library of PAL^37^. After pre-screening with three rounds of growth enrichment, we transformed the plasmids into the biosensor strain, Ecvdt46. We noted that >80 % of the passage #3 population had higher fluorescence than parental PAL (**Figure 4A**), further validating the utility of the pre-screen. We sorted the library based on fluorescence and collected 80 events from top 5 % of the population in a microtiter plate and them screened for tCA production and fluorescence in a spectrophotometer (**Figure 4B-C**). Of these, 11 variants did not revive. The surviving variants showed good correlation (Pearson correlation coefficient, rP = 0.73) between total cellular PAL activity and biosensor output (**Figure 4D**). Of these, we picked the top 14 tCA producers for further characterization and purified them (along with parental PAL) and determined their specific activity in the presence of 30 mM Phe. We observed that biosensor response showed better correlation with PAL specific activity than did growth rate in minimal, ammonium-depleted, Phe medium (**Figure 4E-F**). This suggest that biosensor-based screen is a better to isolate high activity PAL variants compared to the growth-based pre-screen. These results also highlighting the benefit of the dual approach for directed evolution of PAL where the growth-based pre-screen eliminates inactive variants while the biosensor screen stratifies the remaining library by activity. On sequencing these 14 variants we found 20 unique mutated positions (**Table S2**).

**Figure 4.**
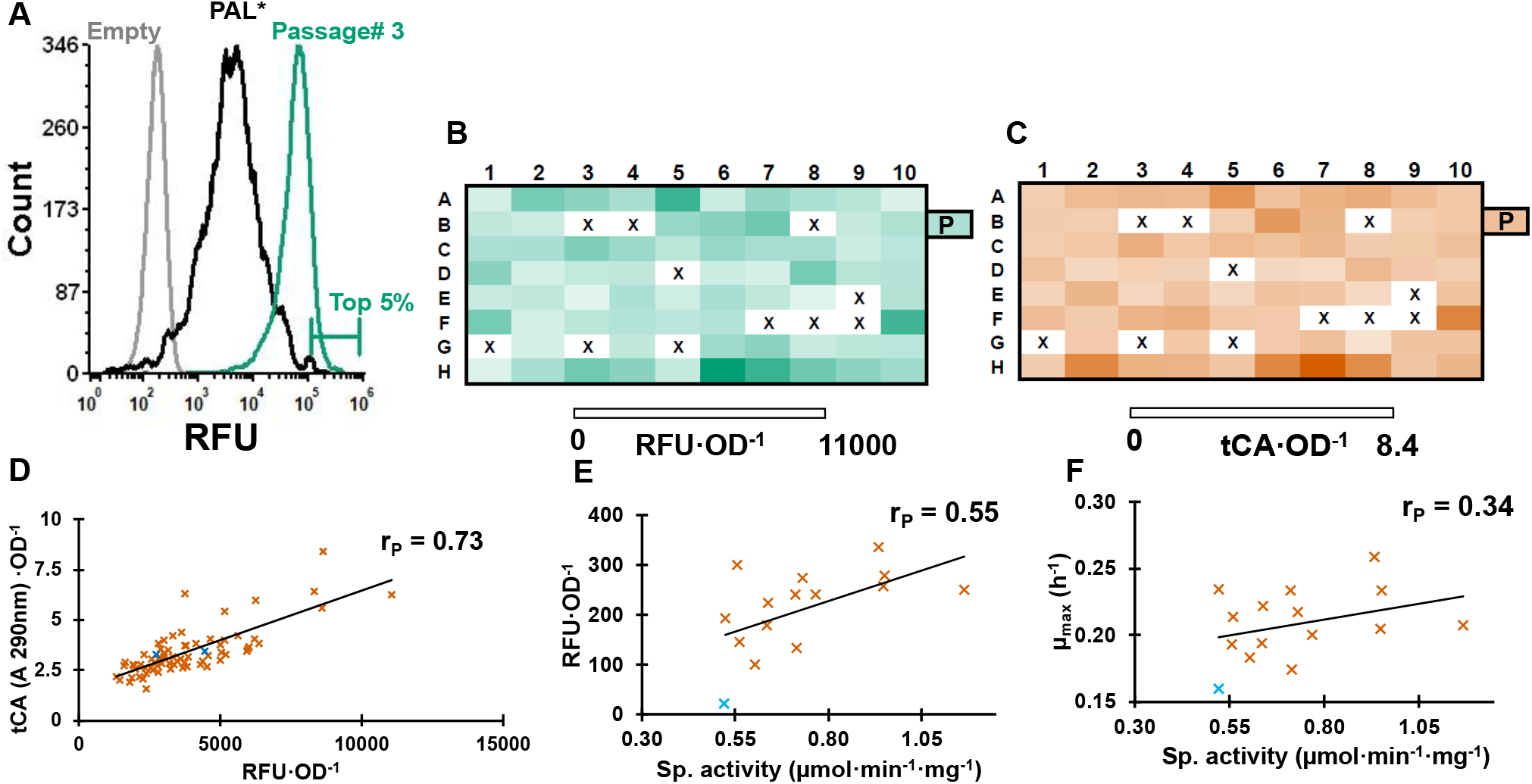
Screening of growth enriched PAL library. **A)** Flow cytometry profile of pre-screened library pool transformed into the biosensor strain. **B)** Biosensor output (RFU·OD^-1^) and **C)** whole cell PAL activity (tCA·OD^-1^) of the top 80 variants collected after FACS of pre-screened pool. P = parental enzyme. x = variants that did not revive. Correlation of the top 14 variants between **D)** cellular tCA production and RFU·OD^-1^, **E)** RFU·OD^-1^ and specific activity of purified variants, and **F)** growth rate in minimal ammonium-depleted Phe-supplemented pre-screen medium and specific activity of purified variants. Mutants are in brown whereas wildtype is blue.

### Further PAL engineering identifies mutants with improved activity

Using these 20 unique positions we created a new library of ~10^7^ members to identify higher activity variants. To achieve this, we generated a new library by combining 3 or 4 mutations/protein (^20^C3-4, 20 choose up to 3 or 4). This strategy was chosen for diversification to ensure that the library size was small enough to be thoroughly screened while also ensuring comprehensive coverage of all 20 positions and 20 amino acids. In this combinatorial approach, we first performed individual site saturation mutagenesis (SSM) at all the twenty positions as separate reactions. Following assembly into circular plasmids and pooling in equimolar amounts, they served as template for second round of individual amplifications using the same SSM primers. Since some of the positions were close together (*viz.* G218, M222), a single SSM primer was generated to span the spatially close sites. Hence, at this stage each variant contained mutations at two to three out of the twenty positions (^20^C2-3). This step was repeated once more to obtain three to four mutations out of the twenty positions (^20^C3-4). This naïve pool was pre-screened and then transformed into the biosensor strain for FACS (**Figure 5A**). Passage #3 of the pre-screen showed higher median sfGFP fluorescence compared to the naïve library, indicating the presence of more active PAL variants. We collected ~10^5^ cells from the top 2 % of the population from both the library pools and screened ~96 variants from each for fluorescence (using a spectrophotometer) and tCA production (**Figure 5B**). Variants from both sorts, naïve and pre-screened, showed good correlation between fluorescence and tCA production. Some variants from the naïve library sort showed low activity indicating continued presence of cheaters. Conversely, all the variants from pre-screened and sorted pool displayed similar or higher activity compared to the parental enzyme. Further, no variants from this pre-screened library were inactive – again, emphasizing the benefit of employing a screening methodology that minimizes cheater enrichment.

**Figure 5.**
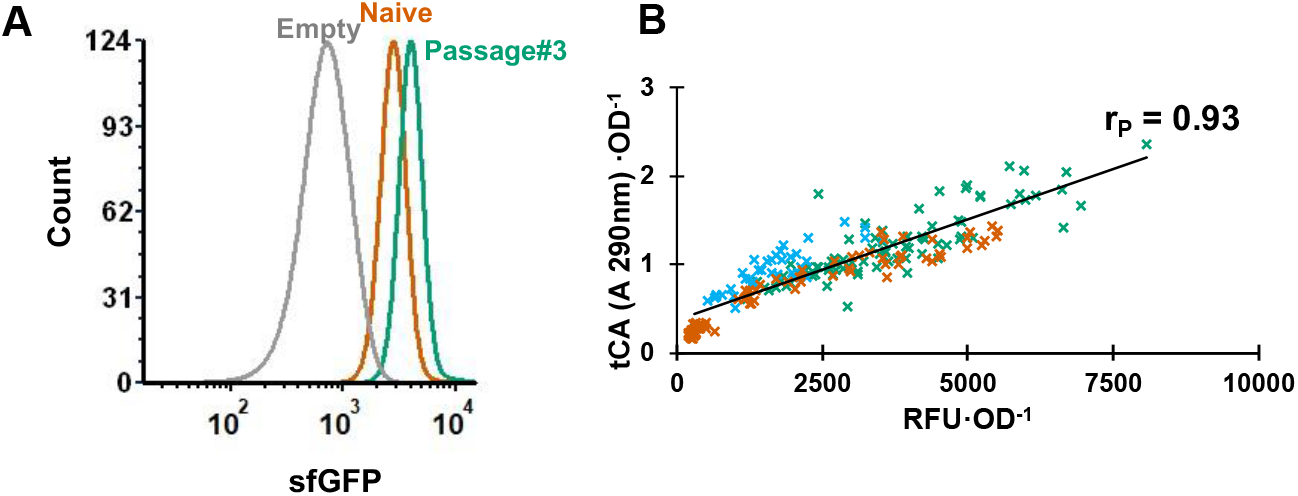
Screening and identification of mutants from a second-generation library. **A)** Flow cytometry profile for Ecvdt46 cells carrying empty plasmid (gray), naïve library (orange) and growth-enriched passage #3 library (green). Refer to supplementary **Figure S4** for FACS scatter plots. **B)** Correlation between whole cell PAL activity (tCA·OD^-1^) and sfGFP fluorescence (RFU·OD^-1^) for variants isolated after FACS of naïve (orange) and pre-screened (green) libraries. Parental PAL is represented in blue, rP represents Pearson correlation coefficient across all datapoints. The sorted naïve library shows a fraction of inactive variants whereas the pre-screened library is largely comprised of variants with higher activity relative to parent.

We sequenced the top 11 variants, performed the Michaelis-Menten kinetic characterization using purified enzymes, and determined their whole cell tCA conversion (**Figure 6A-B**). Of the sequenced 11 variants, S98Y-T102K-L566G was represented thrice. We did not detect any M222L or G218S variants, which were also well-represented in the hits from the first library (**Table S2**) and also identified previously^37^. Thus, in all, we identified 8 unique variants characterized them to determine their kinetic constants (**Figure 6A, Table 1**). All the variants displayed typical Michaelis-Menten behavior with S98Y-T102K-L566G showing the highest increase in k_cat_ (~1.7-fold). Except for two variants, S98T-T102A and S98L-T102A-L566A, the rest showed improved k_cat_ compared to the parental enzyme. Correspondingly, we observed the variants with improved activity to exhibit higher whole cell tCA production (**Figure 6B**). Notably, most of the variants identified in the current screen had mutations at S98, T102, and L566. Of these, only T012 has previously been shown to enhance PAL activity^43^. Further, these positions showed permissivity to biochemically diverse amino acids. For instance, we observed Tyr, Val, Leu, Thr, and Arg at S98, Ala, Ser, and Lys at T102, and Ala, Gly, Ser, Thr, and Glu at L566. We mapped the sampled positions onto the PAL structure (PDB: 3CZO) to gain insights into their functional significance (**Figure S5**). We found that S98 and T102 surround the active site pocket of PAL, whereas L566 is present in the region that is not well defined in the crystal structure. Though residues S98 and T102 have not been shown to be catalytically important, it would be interesting to understand their functional significance in future studies.

**Figure 6.**
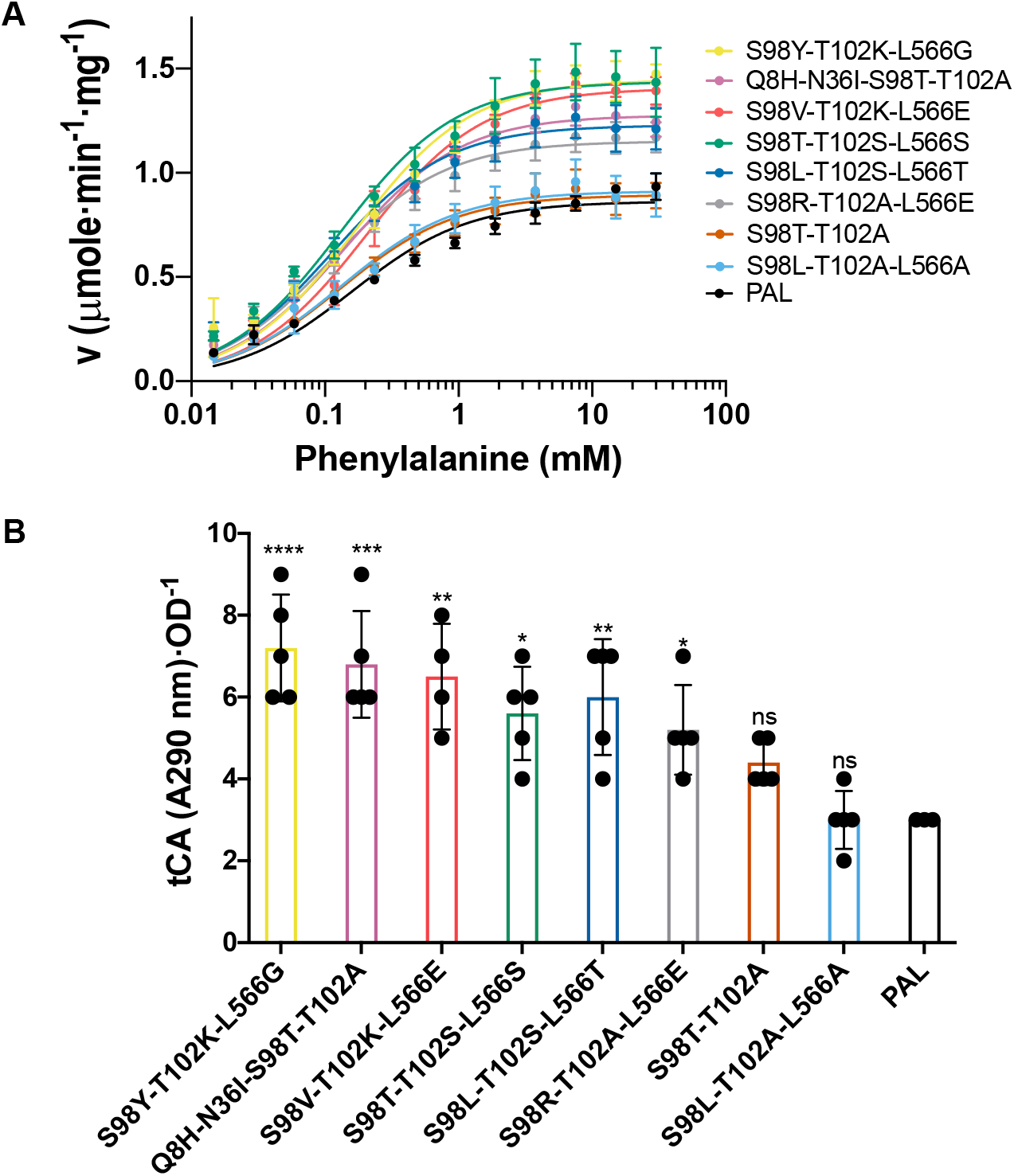
Characterization of variants from passage #3 of second-generation library. **A)** Michaelis-Menten profile of variants. **B)** Whole cell tCA production by PAL variants. **** (p < 0.0001), *** (p ≤ 0.003), ** (p ≤ 0.001), * (p ≤ 0.05), ns (non-significant).

**Table 1.** Kinetic constants for PAL variants.

| Variant | $V_{max}$<br>($\mu\text{mole} \cdot \text{min}^{-1} \cdot \text{mg}^{-1}$) | $K_M$<br>($\mu\text{M}$) | $k_{cat}$<br>($\text{s}^{-1}$) | $k_{cat}/K_M$<br>($\text{s}^{-1} \cdot \text{M}^{-1}$) | Fold increase<br>$k_{cat}$ |
| --- | --- | --- | --- | --- | --- |
| PAL (parental) | $0.86 \pm 0.02$ | $160 \pm 17$ | 0.90 | 5.6 | 1.00 |
| S98Y-T102K-L566G | $1.45 \pm 0.02$ | $168 \pm 14$ | 1.51 | 9.0 | 1.68 |
| Q8H-N36I-S98T-T102A | $1.28 \pm 0.01$ | $141 \pm 08$ | 1.33 | 9.4 | 1.48 |
| S98V-T102K-L566E | $1.41 \pm 0.02$ | $205 \pm 16$ | 1.46 | 7.2 | 1.63 |
| S98T-T102S-L566S | $1.44 \pm 0.03$ | $131 \pm 13$ | 1.50 | 11.4 | 1.67 |
| S98L-T102S-L566T | $1.23 \pm 0.02$ | $112 \pm 10$ | 1.28 | 11.4 | 1.42 |
| S98R-T102A-L566E | $1.15 \pm 0.02$ | $108 \pm 09$ | 1.20 | 11.1 | 1.33 |
| S98T-T102A | $0.89 \pm 0.02$ | $131 \pm 12$ | 0.93 | 7.1 | 1.04 |
| S98L-T102A-L566A | $0.91 \pm 0.02$ | $131 \pm 16$ | 0.95 | 7.2 | 1.06 |

## CONCLUSIONS

Our work outlines a methodology to successfully overcome enrichment of false positives during strain/protein engineering campaigns that utilize biosensors to screen combinatorial libraries. We used the tCA biosensor based on the *E. coli hca* operon (including the tCA-binding HcaR TF, its native promoter, as well as its target promoter, *P_hcaE_*) and the PAL enzyme as a model system. We demonstrate that false positives (or cheaters) enrich due to cross-feeding of tCA – the PAL reaction product and biosensor activator. We hypothesized that since these cheaters likely have lower intracellular tCA concentration, decreasing the sensitivity of the biosensor could, at least partially, mitigate cheater enrichment. We leverage the intrinsic CCR on the *hca* operon by glucose to increase the LOD of the system (decreasing its sensitivity) and demonstrate that this suppresses the relative enrichment of cheaters. While this would also inhibit the catabolism of tCA by *E. coli,* which could have been beneficial in mitigating cross-feeding, we believe that the benefits of desensitizing the biosensor likely outweigh any potential benefit of tCA metabolism. Next, we incorporate our previously developed growth-based enrichment to further repress cheater enrichment by promoting growth of cells carrying active PAL. While the growth-based pre-screen was effective at eliminating inactive PAL variants from the pool, it was only modestly able to link cell growth to tCA productivity. Conversely, we found that the desensitized biosensor itself was better at quantitatively linking PAL activity to the reporter output. Thus, a combination of the two screens was able to optimally identify high activity variants from large, mutagenized libraries. Although the variants isolated here were not as active as those identified via a more thorough deep mutational scanning guided engineering approach^43^, the outcomes are more successful than the previously described approach to engineering PAL activity using the same biosensor ^32^. Further, this work required only a single round of FACS sorting at each step, compared to the previously approach, where five rounds were needed. We expect that a similar analysis and approach could be broadly beneficial to other biosensor-guided engineering workflows that suffer from ligand cross-feeding.

## MATERIALS AND METHODS

### General molecular biology and microbiological techniques

Error-prone PCR on *Anabaena variabilis* PAL was performed as described previously^37^. PCR was performed using Phusion DNA polymerase or Platinum™ SuperFi II Green PCR Master Mix (ThermoFisher Scientific). *E. coli* NEB5α (New England Biolabs) was used for plasmid propagation and *E. coli* MG1655 *rph+* was used for screening of libraries and purification of recombinant PAL and its mutants. Sequences of constructed plasmids were confirmed through DNA sequencing (Genewiz). PAL was expressed under constitutive T5 promoter from plasmid pBAV1k carrying chloramphenicol resistance. BFP-tagged PAL^−^ strain was constructed by knocking-in BFP under IPTG-inducible T7 promoter at *araC* locus using lambda-red recombineering. During the flow cytometry experiments, BFP was induced by adding IPTG (500 μM).

### Enzyme assay, purification, and kinetic characterization

PAL activity was monitored by measuring the production of tCA at 290 nm over time. Briefly, 200 μL reaction as performed by 1 μg of purified enzyme to pre-warmed (37 °C) 1 × PBS containing 30 mM Phe. The assay was performed in 96-well F-bottom UVStar (Greiner Bio-One, Kremsmünster, Austria) microtiter plate and absorbance at 290 nm was measured every 15 s at 37 °C using a SpectraMax M3 (Molecular Devices) plate reader.

The enzyme was purified from 25 mL culture. The pellet was washed once with 1 × PBS and resuspended in 500 μL of the same. This cell suspension was sonicated on ice using a Sonifier SFX 150 (Branson Ultrasonics, Danbury, CT) (10 s ON; 1 min OFF; 2 min; 40 %), and cell debris was separated from the lysate by centrifuging at 20,000 × g for 10 min at 4 °C. As each construct included a N-term His-tag, the enzyme was purified via immobilized metal affinity chromatography (IMAC) purification. Briefly, the lysate was loaded onto HisPur™ Ni-NTA Spin Plates (ThermoFisher Scientific) and incubated for 2 min. After being washed five times with equilibration buffer, pure protein was then eluted using 200 μL of Elution buffer (300 mM NaCl, 50 mM NaH_2_PO_4_, 500 mM imidazole, pH 8.0). Elution fractions were then dialyzed using Tube-O-Dialyser tubes (1 kDa MWCO, Geno-Tech). Protein concentration was estimated by Bradford reagent (VWR) using bovine serum albumin (BSA) as the standard. For kinetic analysis, PALs were purified and assayed as described above. The activity was measured at twelve concentrations of Phe ranging from 15 μM to 30 mM in PBS at 37 °C. A Michaelis-Menten curve was fit in GraphPad Prism software using the initial rate at each Phe concentration.

### Flow cytometry

Flow cytometry analysis was performed using Attune NxT flow cytometer. Relevant flow cytometer settings: 12.5 μL/min flow rate, 20,000 events collected per sample, FSC:120, SSC:240, BL1: 300V, VL1: 250 V. The FCS files were analyzed using FSC software v6.

### Microscopy

Microscopy was performed using DMi8 automated inverted microscope (Leica Microsystems, #11889113) equipped with a CCD camera (Leica Microsystems, #DFC300 G), and a LED405 (Ex 375-435 nm, Em 450-490 nm, exposure time – 10 ms, gain 2) and YFP (Ex 490-510 nm, Em 520-550 nm, exposure time – 5ms, gain 1) filter cube. 1 μL of cultures were spotted on agarose pads (2 % w/v, 1 mm thick).

### Fluorescence-activated cell sorting (FACS)

Cell sorting was performed on a Bio-Rad S3e Cell Sorter. Sorting gates were drawn on dot plots with FL1 and FL4 on the axes. Sorting was performed on the events in the FL1 green emission channel. After sorting, some of the cells were plated on LB + Cm (25 μg·ml^-1^) + Amp (100 μg·ml^-1^) agar plates and remainder was recovered in LB + Cm + Amp liquid medium and frozen at −80 °C for long-term storage.

## ACKNOWLEDGMENTS

We would like to thank all the Nair lab members for helpful comments and insights, and especially Rebecca Condruti for providing inspiration for the title and feedback on the draft. We also like to thank Pramod K. Jangir for providing feedback on the manuscript.

## REFERENCES CITED

1. Rogers, J. K.; Church, G. M., Genetically encoded sensors enable real-time observation of metabolite production. Proceedings of the National Academy of Sciences 2016, 113 (9), 2388–2393.

2. Raman, S.; Rogers, J. K.; Taylor, N. D.; Church, G. M., Evolution-guided optimization of biosynthetic pathways. Proceedings of the National Academy of Sciences 2014, 111 (50), 17803–17808.

3. Dietrich, J. A.; McKee, A. E.; Keasling, J. D., High-throughput metabolic engineering: advances in small-molecule screening and selection. Annual review of biochemistry 2010, 79, 563–590.

4. Zhang, J.; Jensen, M. K.; Keasling, J. D., Development of biosensors and their application in metabolic engineering. Current opinion in chemical biology 2015, 28, 1–8.

5. Zhang, J.; Barajas, J. F.; Burdu, M.; Ruegg, T. L.; Dias, B.; Keasling, J. D., Development of a transcription factor-based lactam biosensor. ACS synthetic biology 2017, 6 (3), 439–445.

6. Ho, J. C.; Pawar, S. V.; Hallam, S. J.; Yadav, V. G., An improved whole-cell biosensor for the discovery of lignin-transforming enzymes in functional metagenomic screens. ACS synthetic biology 2018, 7 (2), 392–398.

7. de los Santos, E. L.; Meyerowitz, J. T.; Mayo, S. L.; Murray, R. M., Engineering transcriptional regulator effector specificity using computational design and in vitro rapid prototyping: developing a vanillin sensor. ACS synthetic biology 2016, 5 (4), 287–295.

8. Lim, H. G.; Jang, S.; Jang, S.; Seo, S. W.; Jung, G. Y., Design and optimization of genetically encoded biosensors for high-throughput screening of chemicals. Current opinion in biotechnology 2018, 54, 18–25.

9. Wu, Y.; Du, G.; Chen, J.; Liu, L., Genetically encoded biosensors and their applications in the development of microbial cell factories. In Engineering of Microbial Biosynthetic Pathways, Springer: 2020; pp 53–73.

10. Sherwood, A. V.; Henkin, T. M., Riboswitch-mediated gene regulation: novel RNA architectures dictate gene expression responses. Annual review of microbiology 2016, 70, 361–374.

11. Jang, S.; Jang, S.; Xiu, Y.; Kang, T. J.; Lee, S.-H.; Koffas, M. A.; Jung, G. Y., Development of artificial riboswitches for monitoring of naringenin in vivo. ACS synthetic biology 2017, 6 (11), 2077–2085.

12. Xiu, Y.; Jang, S.; Jones, J. A.; Zill, N. A.; Linhardt, R. J.; Yuan, Q.; Jung, G. Y.; Koffas, M. A., Naringenin-responsive riboswitch-based fluorescent biosensor module for Escherichia coli co-cultures. Biotechnology and Bioengineering 2017, 114 (10), 2235–2244.

13. Patel, P., (Bio) sensors for measurement of analytes implicated in food safety: a review. TrAC Trends in Analytical Chemistry 2002, 21 (2), 96–115.

14. Ivnitski, D.; Abdel-Hamid, I.; Atanasov, P.; Wilkins, E.; Stricker, S., Application of electrochemical biosensors for detection of food pathogenic bacteria. Electroanalysis: An International Journal Devoted to Fundamental and Practical Aspects of Electroanalysis 2000, 12 (5), 317–325.

15. Trojanowicz, M., Determination of pesticides using electrochemical enzymatic biosensors. Electroanalysis 2002, 14 (19-20), 1311–1328.

16. Pu, J.; Zinkus-Boltz, J.; Dickinson, B. C., Evolution of a split RNA polymerase as a versatile biosensor platform. Nature chemical biology 2017, 13 (4), 432–438.

17. Meyer, A. J.; Segall-Shapiro, T. H.; Glassey, E.; Zhang, J.; Voigt, C. A., Escherichia coli “Marionette” strains with 12 highly optimized small-molecule sensors. Nature chemical biology 2019, 15 (2), 196–204.

18. David, F.; Nielsen, J.; Siewers, V., Flux control at the malonyl-CoA node through hierarchical dynamic pathway regulation in Saccharomyces cerevisiae. ACS synthetic biology 2016, 5 (3), 224–233.

19. Li, S.; Si, T.; Wang, M.; Zhao, H., Development of a synthetic malonyl-CoA sensor in Saccharomyces cerevisiae for intracellular metabolite monitoring and genetic screening. ACS synthetic biology 2015, 4 (12), 1308–1315.

20. De Paepe, B.; Maertens, J.; Vanholme, B.; De Mey, M., Modularization and response curve engineering of a naringenin-responsive transcriptional biosensor. ACS synthetic biology 2018, 7 (5), 1303–1314.

21. Zhou, S.; Yuan, S.-F.; Nair, P. H.; Alper, H. S.; Deng, Y.; Zhou, J., Development of a growth coupled and multi-layered dynamic regulation network balancing malonyl-CoA node to enhance (2S)-naringenin biosynthesis in Escherichia coli. Metabolic Engineering 2021.

22. Wang, G.; Øzmerih, S. l.; Guerreiro, R.; Meireles, A. C.; Carolas, A.; Milne, N.; Jensen, M. K.; Ferreira, B. S.; Borodina, I., Improvement of cis, cis-muconic acid production in Saccharomyces cerevisiae through biosensor-aided genome engineering. ACS synthetic biology 2020, 9 (3), 634–646.

23. Jensen, E. D.; Ambri, F.; Bendtsen, M. B.; Javanpour, A. A.; Liu, C. C.; Jensen, M. K.; Keasling, J. D., Integrating continuous hypermutation with high-throughput screening for optimization of cis, cis-muconic acid production in yeast. Microbial Biotechnology 2021.

24. Zheng, S.; Hou, J.; Zhou, Y.; Fang, H.; Wang, T.-T.; Liu, F.; Wang, F.-S.; Sheng, J.-Z., One-pot two-strain system based on glucaric acid biosensor for rapid screening of myo-inositol oxygenase mutations and glucaric acid production in recombinant cells. Metabolic engineering 2018, 49, 212–219.

25. Dabirian, Y.; Gonçalves Teixeira, P.; Nielsen, J.; Siewers, V.; David, F., FadR-based biosensor-assisted screening for genes enhancing fatty Acyl-CoA pools in Saccharomyces cerevisiae. ACS synthetic biology 2019, 8 (8), 1788–1800.

26. Kunjapur, A. M.; Prather, K. L., Development of a vanillate biosensor for the vanillin biosynthesis pathway in E. coli. ACS synthetic biology 2019, 8 (9), 1958–1967.

27. Jha, R. K.; Bingen, J. M.; Johnson, C. W.; Kern, T. L.; Khanna, P.; Trettel, D. S.; Strauss, C. E.; Beckham, G. T.; Dale, T., A protocatechuate biosensor for Pseudomonas putida KT2440 via promoter and protein evolution. Metabolic engineering communications 2018, 6, 33–38.

28. Yu, H.; Chen, Z.; Wang, N.; Yu, S.; Yan, Y.; Huo, Y.-X., Engineering transcription factor BmoR for screening butanol overproducers. Metabolic engineering 2019, 56, 28–38.

29. Roth, T. B.; Woolston, B. M.; Stephanopoulos, G.; Liu, D. R., Phage-assisted evolution of Bacillus methanolicus methanol dehydrogenase 2. ACS synthetic biology 2019, 8 (4), 796–806.

30. Popa, S. C.; Inamoto, I.; Thuronyi, B. W.; Shin, J. A., Phage-Assisted Continuous Evolution (PACE): A Guide Focused on Evolving Protein–DNA Interactions. ACS omega 2020, 5 (42), 26957–26966.

31. Inamoto, I.; Sheoran, I.; Popa, S. C.; Hussain, M.; Shin, J. A., Combining Rational Design and Continuous Evolution on Minimalist Proteins That Target the E-box DNA Site. ACS Chemical Biology 2020, 16 (1), 35–44.

32. Flachbart, L. K.; Sokolowsky, S.; Marienhagen, J., Displaced by deceivers: prevention of biosensor cross-talk is pivotal for successful biosensor-based high-throughput screening campaigns. ACS synthetic biology 2019, 8 (8), 1847–1857.

33. van Sint Fiet, S.; van Beilen, J. B.; Witholt, B., Selection of biocatalysts for chemical synthesis. Proceedings of the National Academy of Sciences 2006, 103 (6), 1693–1698.

34. Dietrich, J. A.; Shis, D. L.; Alikhani, A.; Keasling, J. D., Transcription factor-based screens and synthetic selections for microbial small-molecule biosynthesis. ACS synthetic biology 2013, 2 (1), 47–58.

35. Woolston, B. M.; Roth, T.; Kohale, I.; Liu, D. R.; Stephanopoulos, G., Development of a formaldehyde biosensor with application to synthetic methylotrophy. Biotechnology and bioengineering 2018, 115 (1), 206–215.

36. Le, T.-K.; Ju, S.-B.; Lee, H.-W.; Lee, J.-Y.; Oh, S.-H.; Kwon, K.-K.; Sung, B.-H.; Lee, S.-G.; Yeom, S.-J., Biosensor-Based Directed Evolution of Methanol Dehydrogenase from Lysinibacillus xylanilyticus. International Journal of Molecular Sciences 2021, 22 (3), 1471.

37. Mays, Z. J.; Mohan, K.; Trivedi, V. D.; Chappell, T. C.; Nair, N. U., Directed evolution of Anabaena variabilis phenylalanine ammonia-lyase (PAL) identifies mutants with enhanced activities. Chemical Communications 2020, 56 (39), 5255–5258.

38. Díaz, E.; Ferrández, A.; García, J. L., Characterization of the hca cluster encoding the dioxygenolytic pathway for initial catabolism of 3-phenylpropionic acid in Escherichia coli K-12. Journal of bacteriology 1998, 180 (11), 2915–2923.

39. Wang, B.; Barahona, M.; Buck, M., Amplification of small molecule-inducible gene expression via tuning of intracellular receptor densities. Nucleic acids research 2015, 43 (3), 1955–1964.

40. Blazeck, J.; Alper, H. S., Promoter engineering: recent advances in controlling transcription at the most fundamental level. Biotechnology journal 2013, 8 (1), 46–58.

41. Hammer, K.; Mijakovic, I.; Jensen, P. R., Synthetic promoter libraries–tuning of gene expression. Trends in biotechnology 2006, 24 (2), 53–55.

42. Turlin, E.; Perrotte-piquemal, M.; Danchin, A.; Biville, F., Regulation of the early steps of 3-phenylpropionate catabolism in Escherichia coli. Journal of molecular microbiology and biotechnology 2001, 3 (1), 127–133.

43. Trivedi, V. D.; Chappell, T. C.; Krishna, N. B.; Shetty, A.; Sigamani, G. G.; Mohan, K.; Ramesh, A.; Kumar, P.; Nair, N. U., In-depth sequence-function characterization reveals multiple paths to enhance phenylalanine ammonia-lyase (PAL) activity. bioRxiv 2021.

