## Supplemental Information for "Cheating the cheater: Suppressing false positive enrichment during biosensor-guided biocatalyst engineering"

**C**

**B**

**A**

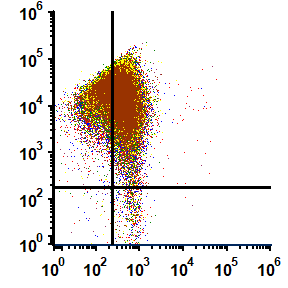

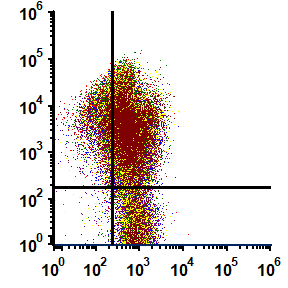

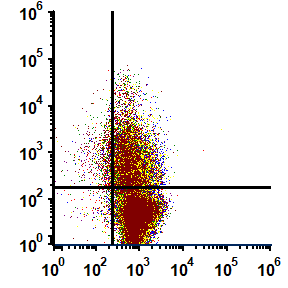

**16%**

**52%**

**65%**

**D**

**E**

**F**

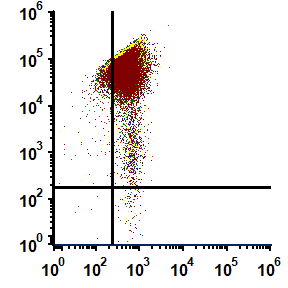

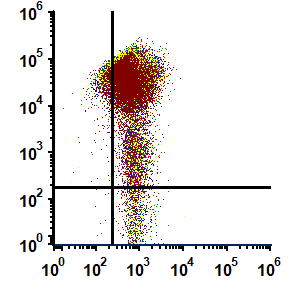

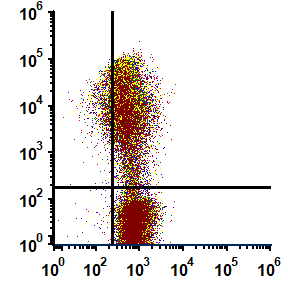

**sfGFP**

**22%**

**65%**

**85%**

**BFP**

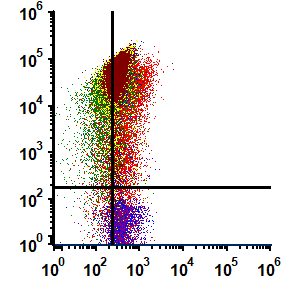

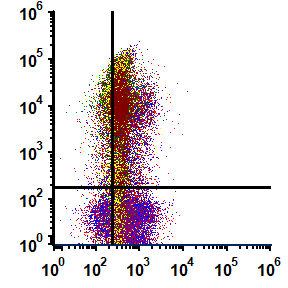

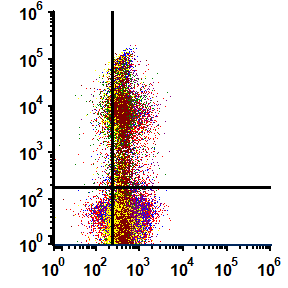

**46%**

**22%**

**13%**

**G**

**H**

**I**

**J**

**K**

**L**

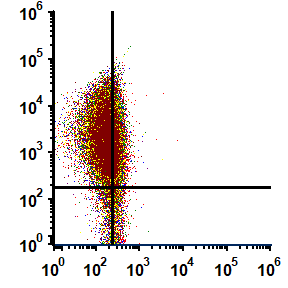

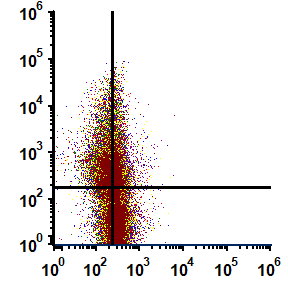

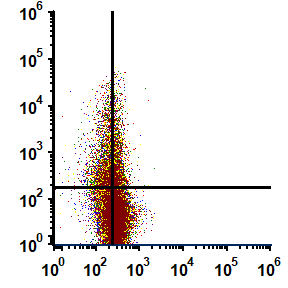

**BFP**

**sfGFP**

**4%**

**15%**

**17%**

**Figure S1**. Flowcytometry profile of Figure 2G. PAL^+^:PAL^–^ in 1:1, 1:10, 1:100 in **A-C)** LB, **D-F)** LB + Phe (30 mM), **G-I)** glycerol + Phe (30 mM), **J-L)** glucose + Phe (30 mM), respectively. Different color scatter plots indicates the data from multiple replicates.

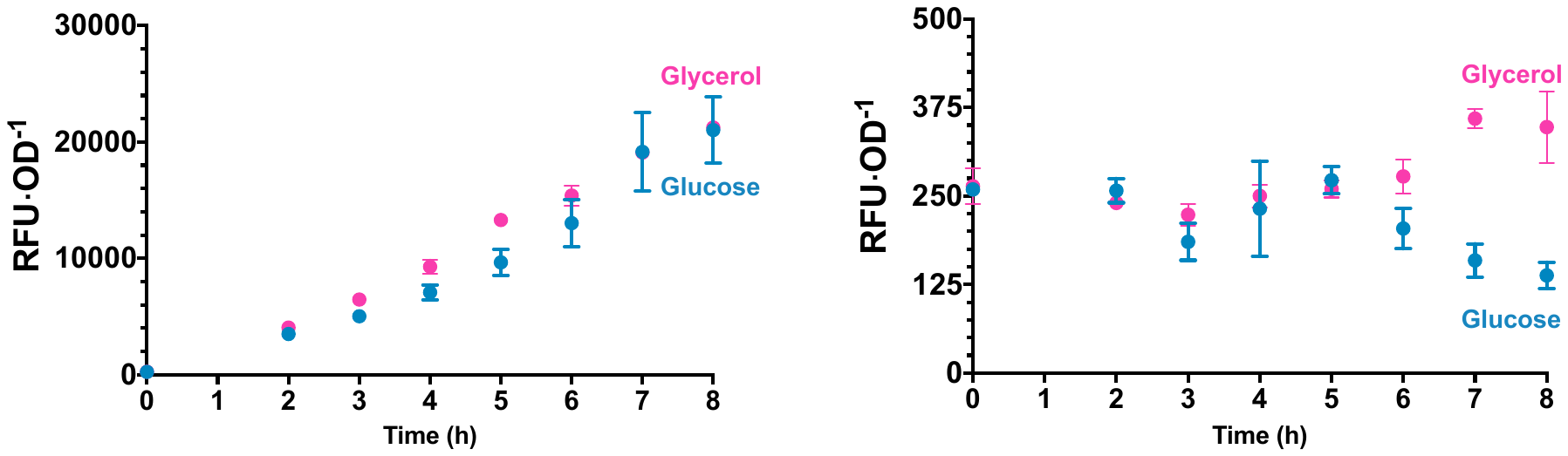

**B**

**A**

**Figure S2**. Response of tCA sensor strain (Ecvdt46) in glycerol (pink) and glucose (blue) containing media in presence of **A)** tCA (0.75 μM) and **B)** without tCA. Though Ecvdt46 response is nearly same in presence of tCA, the presence of glucose suppresses the activation of sensor in absence of tCA, giving better signal:noise.

**D**

**C**

**B**

**A**

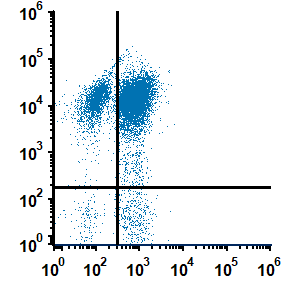

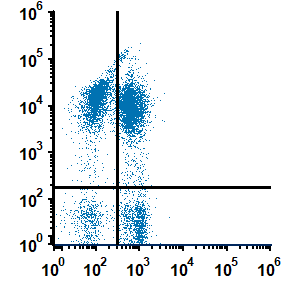

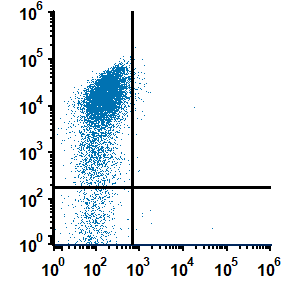

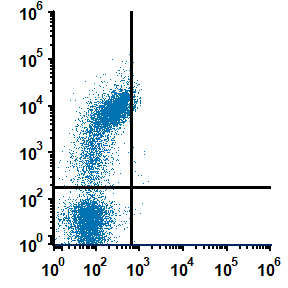

**2%**

**1%**

**42%**

**66%**

**BFP**

**sfGFP**

**Figure S3**. Flow cytometry profile of mock library containing PAL^+^:PAL^–^ in 1:1 ratio and enriched in glucose + Phe (30 mM). **A)** naïve library, **B-D)** passage #1, #2, and #3, respectively. The values in upper right quadrant indicate the percentage of cheater cells.

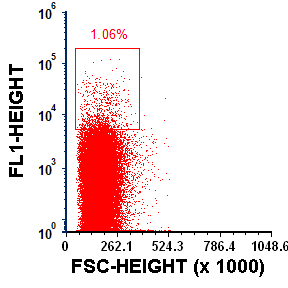

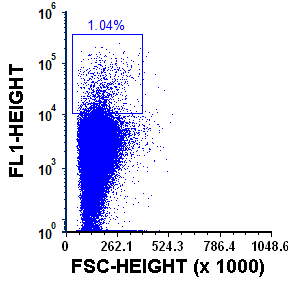

**B**

**A**

**Figure S4**. FACS profile of ^20^C_3−4_ library. **A)** Naïve and **B)** passage #3 post growth screen. Inset plots indicates the top ~1 % of cells that were collected from both the samples for further analysis. FL1-Height indicates the sfGFP fluorescence.

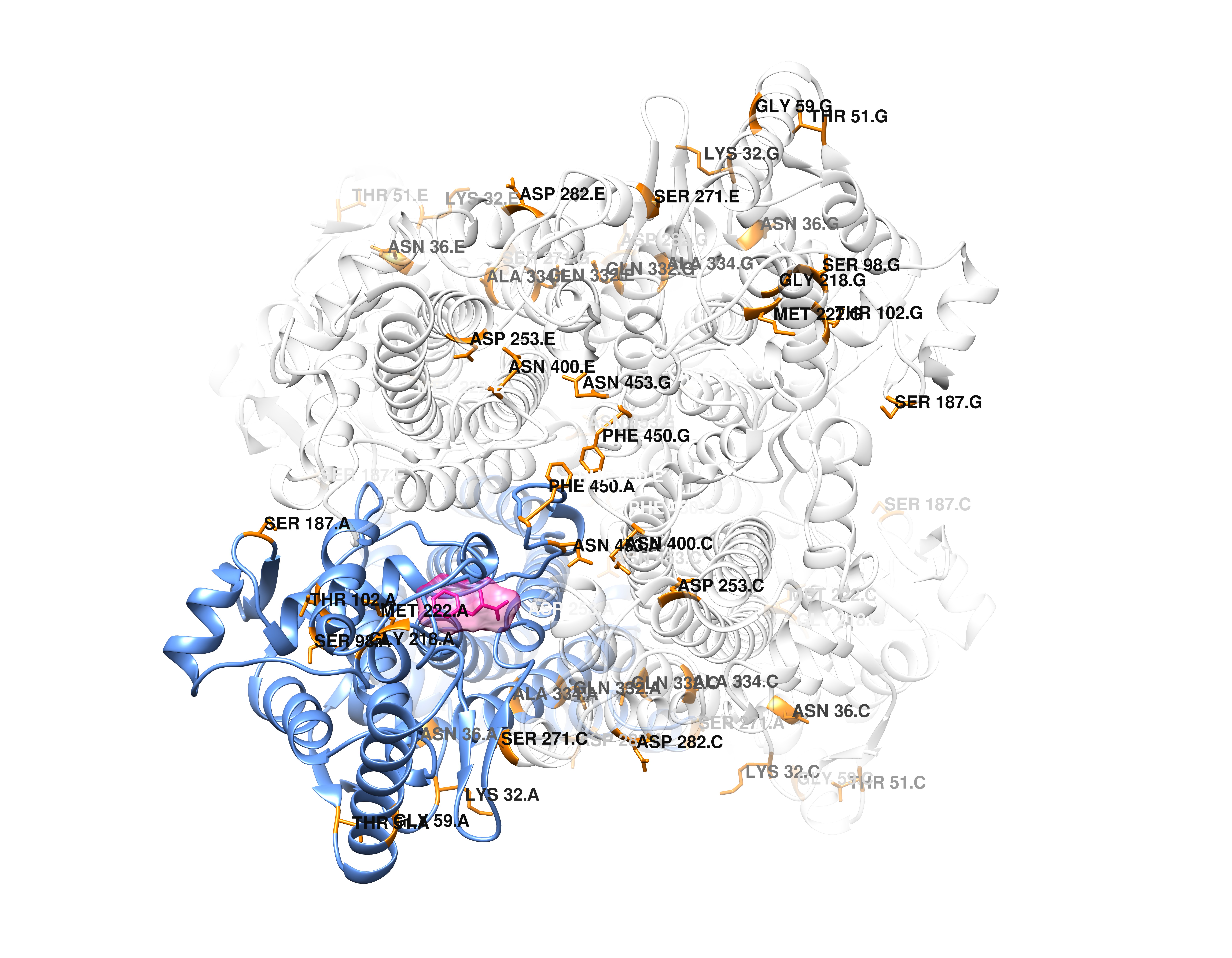

**Figure S5**. Mutants identified in the present study mapped onto the structure of PAL (PDB 3CZO). Subunit A is indicated in blue. The other three subunits are as faint outlines to indicate the oligomerization state of PAL.

**Table S1**. Flow cytometry data of PAL^+^:PAL^–^ in 1:1, 1:10, 1:100 in LB, LB + Phe (30 mM), glycerol + Phe (30 mM), glucose + Phe (30 mM)

|  |  |  | **LB** | | | **LB + Phe (30 mM )** | | | **Gly + Phe (30 mM)** | | | **Glc + Phe (30 mM)** | | |
| --- | --- | --- | --- | --- | --- | --- | --- | --- | --- | --- | --- | --- | --- | --- |
| **Replicates** |  | **Gate** | 1:1 | 1:10 | 1:100 | 1:1 | 1:10 | 1:100 | 1:1 | 1:10 | 1:100 | 1:1 | 1:10 | 1:100 |
| **1** |  | UL | 20 | 10 | 1 | 5 | 3 | 2 | 13 | 4 | 2 | 40 | 16 | 6 |
|  | Cheater | UR | 65 | 55 | 13 | 88 | 66 | 24 | 58 | 22 | 13 | 22 | 13 | 5 |
|  |  | LL | 1 | 0 | 0 | 0 | 0 | 0 | 8 | 14 | 11 | 11 | 17 | 22 |
|  |  | LR | 14 | 34 | 85 | 7 | 31 | 74 | 22 | 60 | 74 | 27 | 53 | 67 |
| **2** |  | UL | 21 | 8 | 1 | 4 | 3 | 2 | 5 | 3 | 1 | 43 | 16 | 4 |
|  | Cheater | UR | 64 | 50 | 16 | 89 | 66 | 22 | 38 | 19 | 6 | 16 | 14 | 3 |
|  |  | LL | 1 | 0 | 0 | 0 | 0 | 0 | 5 | 11 | 5 | 18 | 16 | 27 |
|  |  | LR | 14 | 42 | 83 | 7 | 31 | 76 | 51 | 67 | 87 | 23 | 54 | 66 |
| **3** |  | UL | 23 | 7 | 1 | 4 | 3 | 2 | 8 | 3 | 1 | 42 | 16 | 4 |
|  | Cheater | UR | 63 | 50 | 14 | 89 | 65 | 21 | 48 | 21 | 13 | 16 | 22 | 3 |
|  |  | LL | 1 | 0 | 0 | 0 | 0 | 0 | 4 | 9 | 4 | 18 | 10 | 32 |
|  |  | LR | 13 | 42 | 85 | 6 | 32 | 76 | 40 | 66 | 82 | 24 | 53 | 61 |
| **4** |  | UL | 23 | 8 | 2 | 5 | 3 | 2 | 28 | 7 | 3 | 45 | 19 | 7 |
|  | Cheater | UR | 63 | 52 | 15 | 86 | 64 | 22 | 32 | 27 | 20 | 19 | 13 | 6 |
|  |  | LL | 1 | 0 | 0 | 0 | 0 | 0 | 28 | 13 | 9 | 16 | 19 | 23 |
|  |  | LR | 12 | 39 | 83 | 8 | 32 | 76 | 12 | 54 | 68 | 20 | 48 | 65 |
| **5** |  | UL | 22 | 7 | 1 | 4 | 3 | 2 | 15 | 4 | 2 | 44 | 16 | 6 |
|  | Cheater | UR | 65 | 51 | 19 | 89 | 64 | 23 | 50 | 18 | 8 | 16 | 15 | 5 |
|  |  | LL | 1 | 1 | 0 | 1 | 0 | 0 | 11 | 14 | 9 | 18 | 17 | 22 |
|  |  | LR | 12 | 41 | 80 | 7 | 33 | 75 | 24 | 64 | 81 | 23 | 52 | 67 |
| **6** |  | UL | 21 | 8 | 2 | 4 | 3 | 2 | 11 | 4 | 1 | 47 | 17 | 7 |
|  | Cheater | UR | 69 | 57 | 18 | 85 | 65 | 23 | 49 | 29 | 17 | 15 | 15 | 6 |
|  |  | LL | 1 | 0 | 0 | 0 | 0 | 0 | 8 | 6 | 2 | 19 | 17 | 23 |
|  |  | LR | 10 | 35 | 79 | 10 | 32 | 75 | 32 | 62 | 80 | 18 | 51 | 65 |

**Table S2**. Top mutants isolated from the epPCR library after pre-screen and a single round of FACS.

| **Variant ID** | **Mutation** |
| --- | --- |
| 1 | M222L |
| 2 | N36S-G218S-I268I |
| 3 | S98T |
| 4 | S98N |
| 5 | G59V |
| 6 | A7S-Q8L-T102S-M222L-L257L-S291S |
| 7 | T46T-T229T-D282E-K301K-Q332K-A334T-N453S |
| 8 | K32E-V556V |
| 9 | M222L-V388V-G483G-L566V |
| 10 | M222L-D253G-D328D-V352V-R442R |
| 11 | T51S-S271G-F450S |
| 12 | T102S-M222L-L257L-S291S |
| 13 | G74G |
| 14 | S187C-N400S |
